# Plant and Bacterial Metaxin-like Proteins: Novel Proteins Related to Vertebrate Metaxins Involved in Uptake of Nascent Proteins into Mitochondria

**DOI:** 10.1101/2020.03.02.972976

**Authors:** Kenneth W. Adolph

## Abstract

The metaxins were originally identified as vertebrate proteins of the outer mitochondrial membrane involved in protein import into mitochondria. Metaxin proteins have also been found in diverse invertebrate phyla. The present study is concerned with examining whether metaxin-like proteins occur in plants and bacteria. Metaxin-like proteins were revealed by their homology with human metaxins and the possession of characteristic GST_Metaxin protein domains. The results demonstrate that metaxin-like proteins exist in plants that include a wide variety of angiosperms, both eudicots and monocots, and other plant groups. Metaxin-like proteins can also be detected in bacteria, particularly in the Proteobacteria phylum, but also in different bacterial phyla. Phylogenetic analysis indicates that plant metaxin-like proteins, bacterial metaxin-like proteins, and vertebrate metaxins form distinct phylogenetic groups, but are related. Metaxin-like proteins, however, are only distantly related to GSTs (glutathione S-transferase proteins). A similar degree of homology is found in aligning the amino acid sequences of plant and bacterial metaxin-like proteins with human metaxins 1, 2, and 3 and other vertebrate metaxins. The amino acid identities range from about 22%-28% for each alignment. The presence of two conserved protein domains, GST_N_Metaxin and GST_C_Metaxin, in both plant and bacterial metaxin-like proteins provides evidence that these proteins are related to the vertebrate and invertebrate metaxins. The metaxin-like proteins have predicted secondary structures that are dominated by alpha-helical segments, like the vertebrate and invertebrate metaxins.

## 1. INTRODUCTION

The vertebrate metaxins, particularly metaxins 1 and 2, are well-established as mitochondrial membrane proteins involved in the uptake of nascent beta-barrel proteins and their incorporation into the outer mitochondrial membrane. In addition, a metaxin 3 gene was identified in the zebrafish *Danio rerio* and in the frog *Xenopus laevis*, and metaxin 3 proteins were found to be highly conserved among vertebrates (Adolph, 2019). Furthermore, metaxin proteins were detected among invertebrates such as the nematode *C. elegans* and the fruit fly *Drosophila melanogaster*. A broad range of invertebrate phyla were found to possess proteins homologous to vertebrate metaxins 1 and 2 (Adolph, 2020). The aim of the study reported here was to investigate whether proteins homologous to the metaxins also exist in plants and bacteria. It was found that metaxin-like proteins are present in a large variety of plant and bacterial species.

For plants and bacteria, deeper understanding of their evolution and functioning is emerging through genome sequencing and analysis. A large number of plant genomes and an even larger number of bacterial genomes have now been completely sequenced. For plants, an important example is *Arabidopsis thaliana*, the most widely used experimental organism in plant biology research. *Arabidopsis* was the first plant to have its genome fully sequenced (The *Arabidopsis* Genome Initiative, 2000). Additional examples that show the variety of plants with sequenced genomes include Asian rice *Oryza sativa* (Ouyang et al., 2007), the poplar tree *Populus trichocarpa* (Tuskan et al., 2006), the grapevine *Vitis vinifera* (The French-Italian Public Consortium for Grapevine Genome Characterization, 2007), and the moss *Physcomitrella patens* (Rensing et al., 2008).

Bacteria with metaxin-like proteins that have had their genomes completely sequenced include *Pseudomonas aeruginosa*, a bacterium associated with hospital-acquired infections (Stover et al., 2000). As other examples, genome sequences have been published for *Francisella tularensis* (Larsson et al., 2005), which causes the disease tularemia, *Acaryochloris marina* (Swingley et al., 2008), a cyanobacterium, *Colwellia psychrerythraea* (Methe et al., 2005), a low temperature and high pressure tolerant bacterium, and *Nitrospira multiformis* (Lucker et al., 2010), which plays a role in the nitrogen cycle.

The mouse metaxin 1 gene (*Mtx1*) was the first metaxin gene to be characterized. Experimental evidence showed that the encoded protein is an outer mitochondrial membrane protein and is involved in the import of proteins into mitochondria (Armstrong et al., 1997). A metaxin 1 gene was also found in humans (*MTX1*), located at cytogenetic location 1q21 between a glucocerebrosidase pseudogene (*psGBA1*) and the gene for thrombospondin 3 (*THBS3*), an extracellular matrix protein (Long et al., 1996; Adolph et al., 1995). A mouse metaxin 2 protein, with 29% amino acid identities compared to mouse metaxin 1, was revealed by its interaction with metaxin 1 protein (Armstrong et al., 1999). It was proposed that a complex consisting of metaxin 1 and metaxin 2 proteins might have a role in the uptake of nascent proteins into mitochondria. A human metaxin 2 gene *(MTX2*) also exists, at 2q31.2.

Metaxin 3 proteins were initially identified through sequencing of cDNAs of zebrafish and *Xenopus* (Adolph, 2004, 2005). Both are classic model vertebrates in biology research. The zebrafish and *Xenopus* metaxin 3 proteins possess the GST_N_Metaxin and GST_C_Metaxin domains also found with vertebrate metaxins 1 and 2. The deduced zebrafish metaxin 3 protein of 313 amino acids shares 40% amino acid identities with zebrafish metaxin 1 and 26% with metaxin 2. For *Xenopus* metaxin 3 aligned with *Xenopus* metaxins 1 and 2, the identities are 39% and 22%, respectively. Zebrafish and *Xenopus* metaxin 3 proteins are 55% homologous.

Protein complexes containing metaxins 1 and 2 in the outer mitochondrial membranes of vertebrates are thought to have equivalent functions to the TOM or SAM complexes of the yeast *Saccharomyces cerevisiae*. Investigations of protein import into yeast mitochondria have revealed detailed information regarding the roles of the TOM and SAM complexes (Pfanner et al., 2019; Wiedemann and Pfanner, 2017; Neupert, 2015). The two complexes work together to bring about the uptake of beta-barrel proteins and other proteins into yeast mitochondria. The TOM37 protein domain, found in the import complexes of yeast, is also a conserved protein domain of vertebrate metaxins and some metaxin-like proteins (this study).

In bacteria, the BamABCD complex is functionally homologous to the SAM complex of *Saccharomyces cerevisiae* and other fungi for the insertion of beta-barrel proteins and other membrane proteins into the bacterial outer membrane. As part of this study, it was found that bacteria containing metaxin-like proteins also contain components of the BamABCD complex. In particular, Bam A is widely distributed among bacteria such as *Pseudomonas* that have metaxin-like proteins. BamB and BamD also show a significant, although lesser, degree of conservation among bacteria with metaxin-like proteins. It cannot therefore be concluded that the metaxin-like proteins are simply substituting for the BamABCD system for insertion of proteins into the bacterial outer membrane.

There have been no publications that report the presence of metaxin-like proteins in plants or bacteria. Because of this, the possible existence of these proteins has been examined. It was found that metaxin-like proteins are present in a wide variety of plants and bacteria, and fundamental aspects of the proteins were investigated.

## 2. MATERIALS AND METHODS

For the phylogenetic analysis of the plant and bacterial metaxin-like proteins shown in Figures 1 and 4, the COBALT multiple sequence alignment tool was used, with phylogenetic trees generated from the alignments (www.ncbi.nlm.nih.gov/tools/cobalt; Papadopoulos and Agarwala, 2007). Alternatively, multiple sequence alignments and phylogenetic trees were produced with Clustal Omega (https://www.ebi.ac.uk/Tools/msa/clustalo/). The Global Align tool of the NCBI (https://blast.ncbi.nlm.nih.gov/Blast.cgi; Needleman and Wunsch, 1970; Altschul et al., 1997) was used to provide the amino acid sequence alignments in Figure 2. Alignments were also generated with the EMBOSS Needle tool of the European Bioinformatics Institute (https://www.ebi.ac.uk/Tools/psa/emboss_needle/). The conserved domain database of the NCBI was searched using the CD_Search tool (www.ncbi.nlm.nih.gov/Structure/cdd/wrpsb.cgi; Marchler-Bauer et al., 2017) to give the domain structures of the metaxin-like proteins included in Figure 3. Determining the genes adjacent to the metaxin-like genes made use of the information on “Gene neighbors” from protein BLAST searches (https://blast.ncbi.nlm.nih.gov/Blast.cgi). Neighboring genes were also revealed by means of the Genome Data Viewer (www.ncbi.nlm.nih.gov/genome/gdv) and, previously, the Map Viewer genome browsers. To predict the alpha-helical and beta-strand secondary structures of the metaxin-like proteins (Figure 5), the PSIPRED secondary structure prediction server was primarily used (bioinf.cs.ucl.ac.uk/psipred/; Jones, 1999). Determining the presence of transmembrane helices employed the PHOBIUS program (www.ebi.ac.uk/Tools/pfa/phobius/; Madeira et al., 2019). Also used was the TMHMM prediction server (www.cbs.dtu.dk/services/TMHMM/; Krogh et al., 2001).

**Figure 1.**
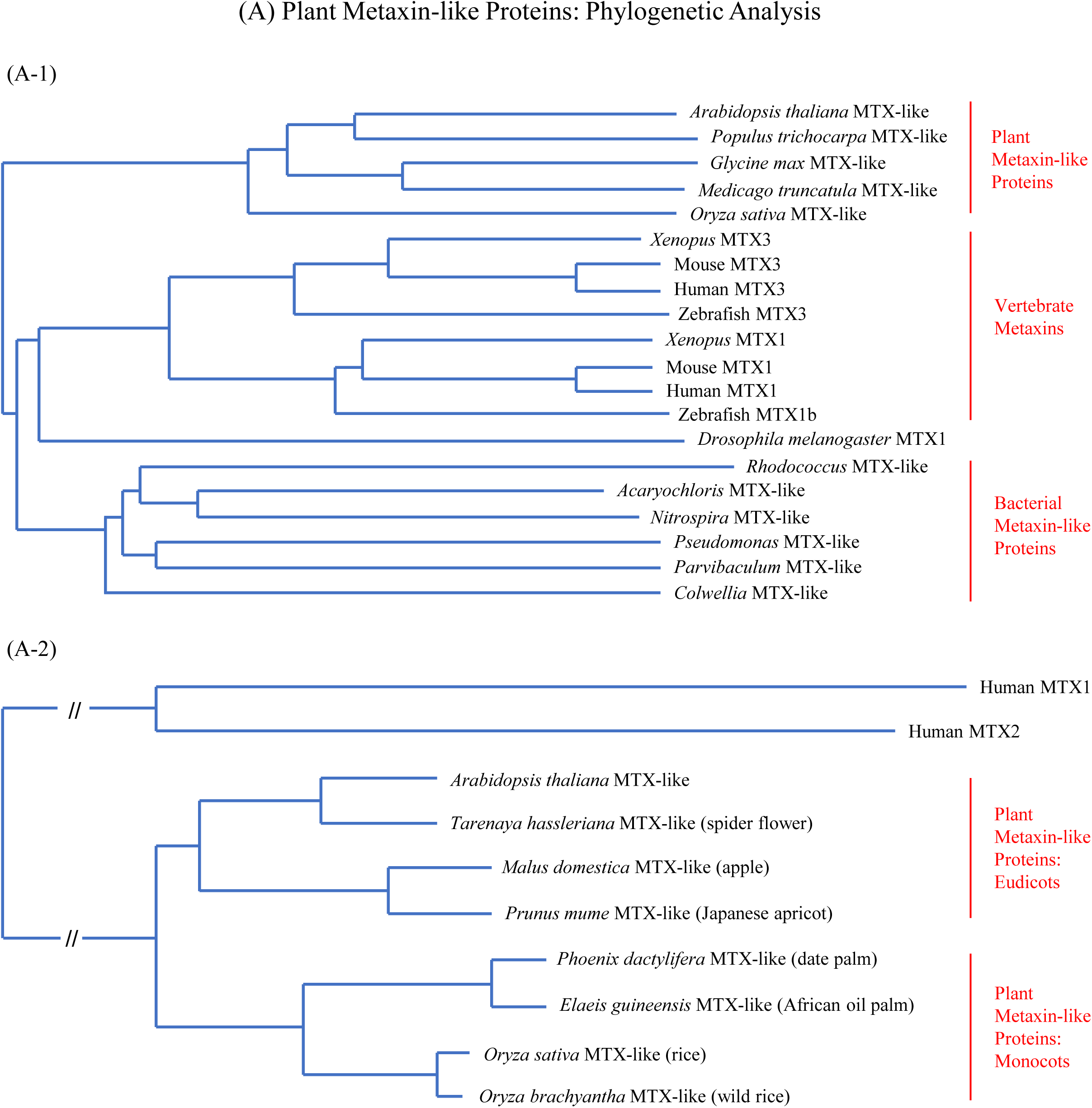
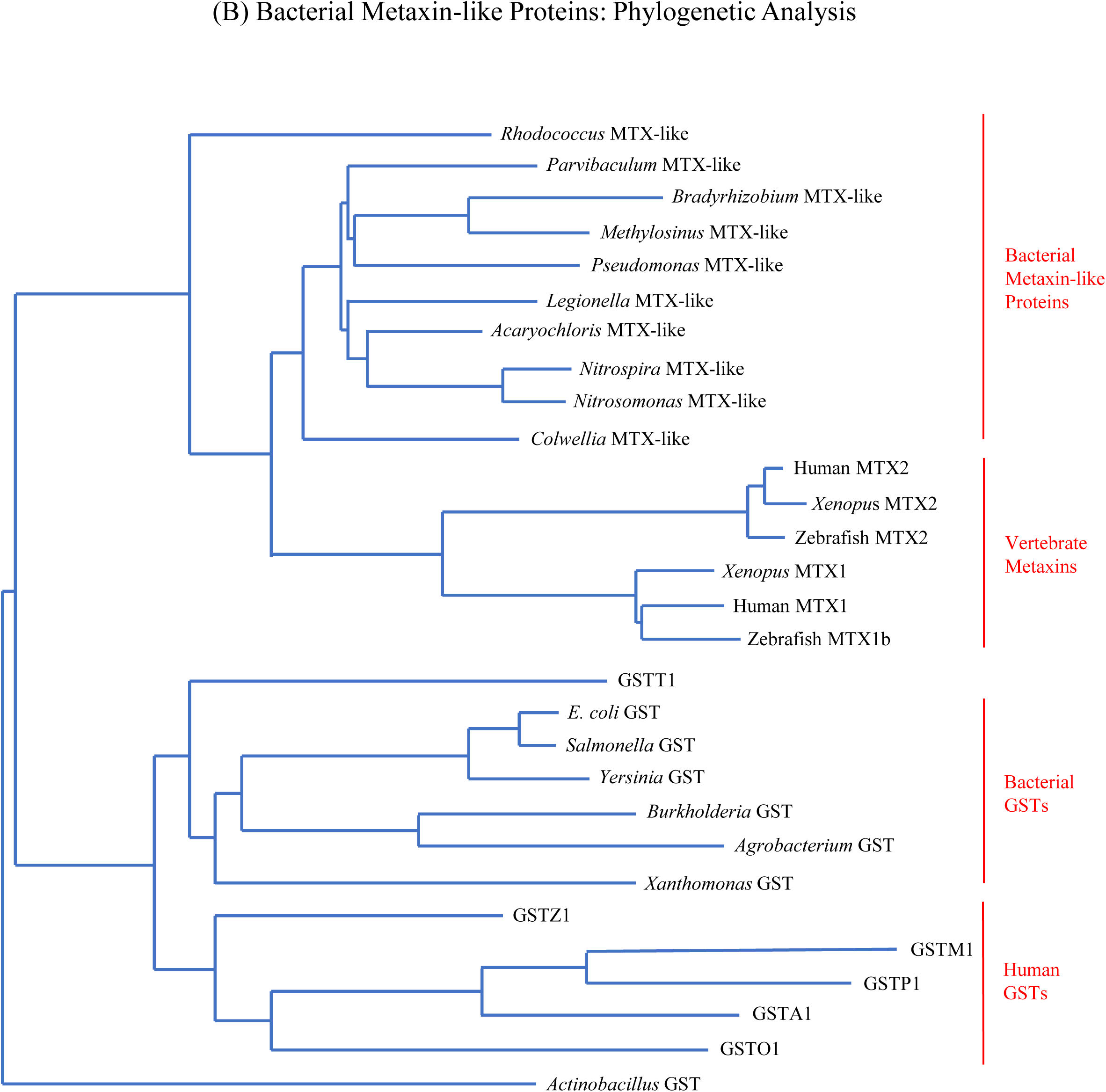
Phylogenetic analysis of plant and bacterial metaxin-like proteins. (A-1) Evolutionary relationships of selected plant metaxin-like proteins, bacterial metaxin-like proteins, and vertebrate metaxins. The plant metaxin-like proteins include that of *Arabidopsis thaliana*, a widely used model organism in plant research. Other plant metaxin-like proteins in Figure 1(A-1) are from the poplar *Populus trichocarpa*, the soybean *Glycine max*, the legume *Medicago truncatula*, and Asian rice *Oryza sativa*. The bacterial metaxin-like proteins in the figure are those of *Rhodococcus* sp. (Phylum: Actinobacteria), *Acaryochloris marina* (Phylum: Cyanobacteria), *Nitrospira multiformis* (Phylum: Nitrospirae), *Pseudomonas aeruginosa* (Phylum: Proteobacteria; Class: Gammaproteobacteria), *Parvibaculum lavamentivorans* (Alphaproteobacteria), and *Colwellia psychrerythraea* (Gammaproteobacteria). Figure 1(A-2) shows that eudicots (*Arabidopsis*, spider flower, apple, apricot) and monocots (date palm, oil palm, rice, wild rice) form separate phylogenetic groups. Also, the metaxin-like proteins cluster together in similar taxonomic groups: *Arabidopsis thaliana* and the cultivated flower *Tarenaya hassleriana*, the apple *Malus domestica* and the apricot *Prunus mume*, the date palm *Phoenix dactylifera* and the oil palm *Elaeis guineensis*, and cultivated rice *Oryza sativa* and *Oryza brachyantha*, a grass in the rice genus. In Figure 1B, the bacterial metaxin-like proteins, in addition to those in Figure 1(A-1), include *Bradyrhizobium* sp. (Phylum: Proteobacteria; Class: Alphaproteobacteria), *Methylosinus trichosporium* (Alphaproteobacteria), *Legionella drancourtii* (Gammaproteobacteria), and *Nitrosomonas* sp. (Betaproteobacteria). The bacterial GSTs (glutathione S-transferase proteins) in Figure 1B are from Gammaproteobacteria (*Escherichia coli* K-12, *Salmonella enterica, Yersinia pestis, Xanthomonas oryzae*, and *Actinobacillus pleuropneumoniae*), Betaproteobacteria (*Burkholderia multivorans*), and Alphaproteobacteria (*Agrobacterium tumefaciens*).

**Figure 2.**
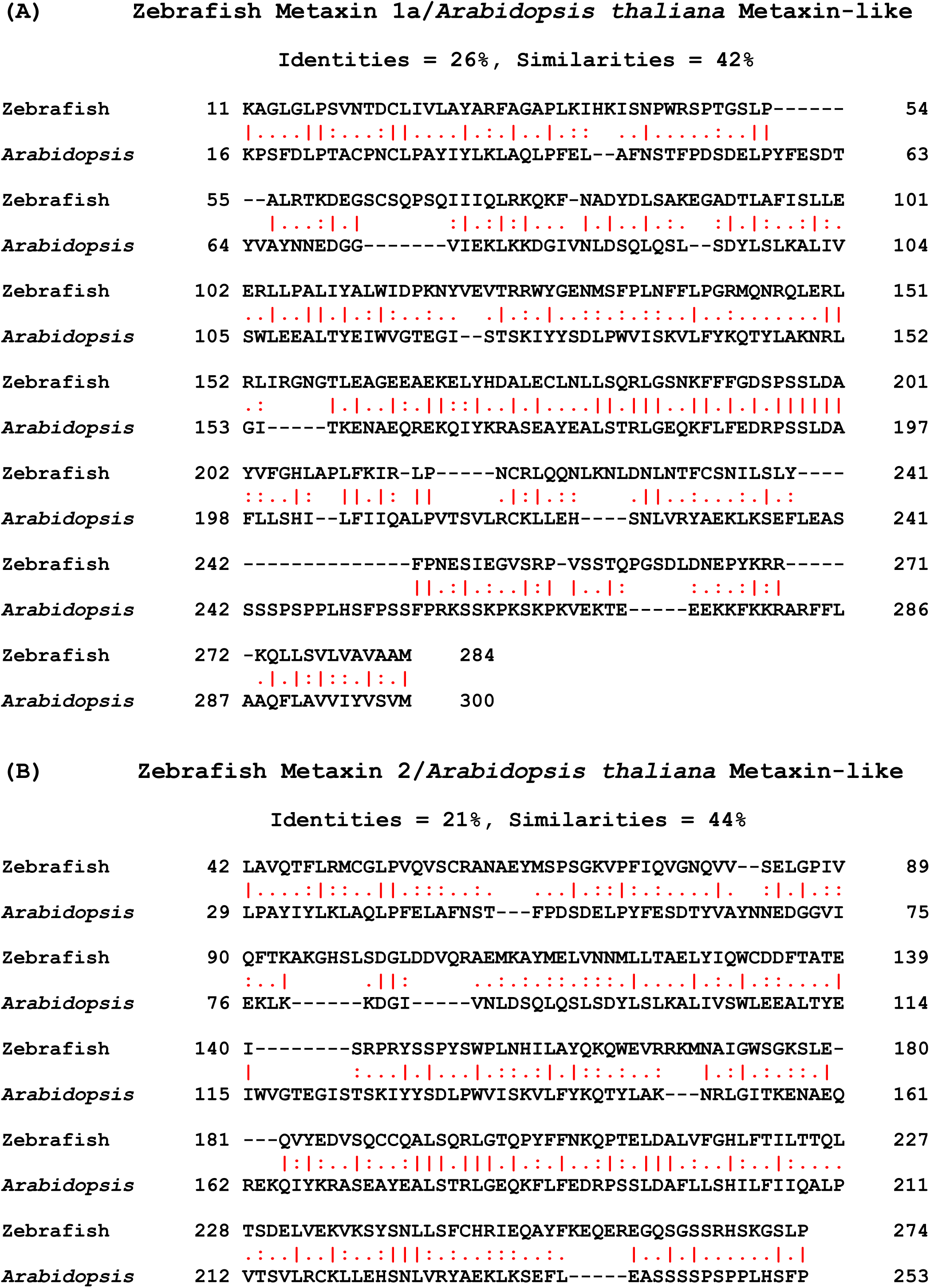
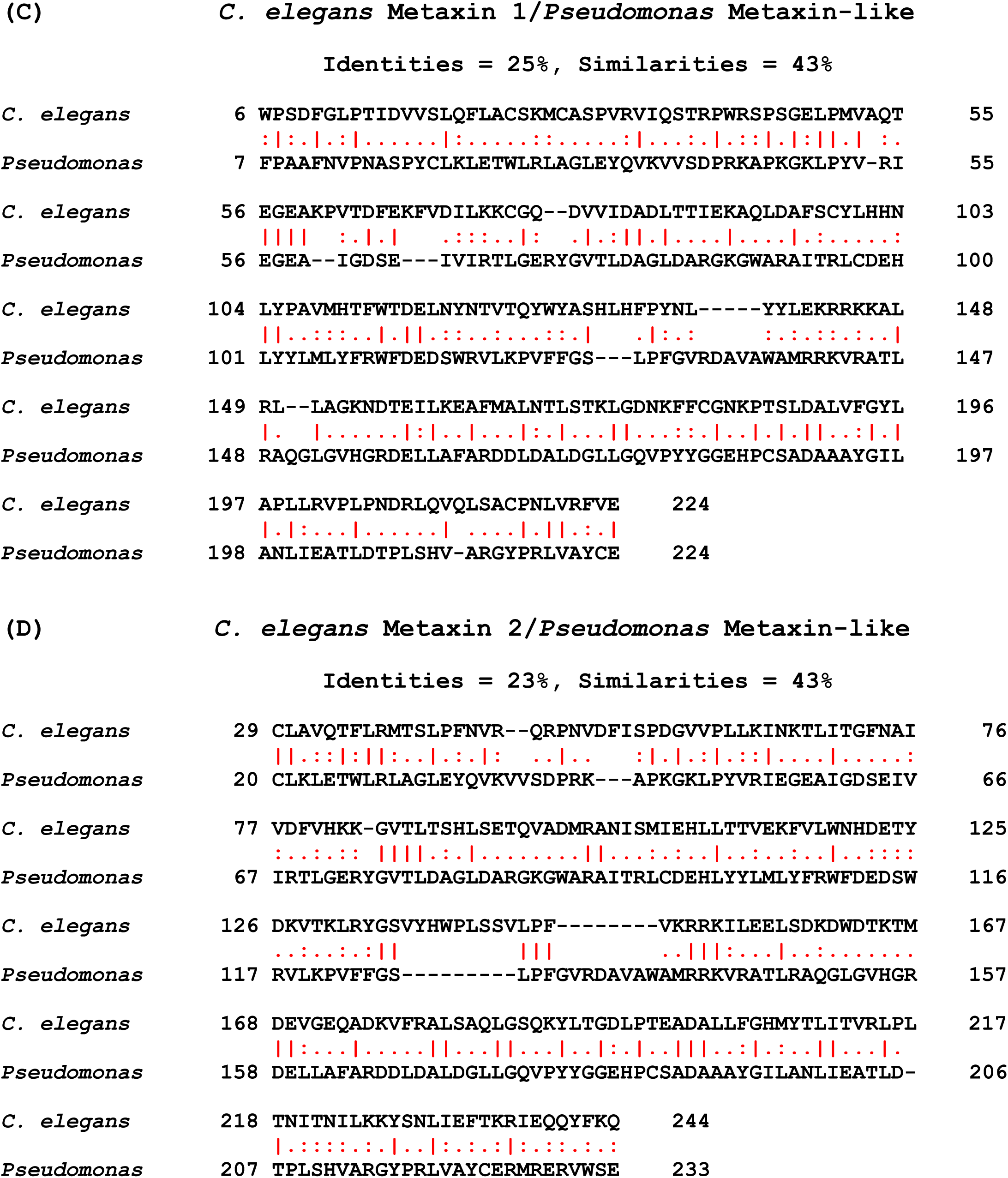
Amino acid sequence identities and similarities of plant and bacterial metaxin-like proteins. (A) The amino acid sequence of zebrafish metaxin 1a is aligned with the sequence of the *Arabidopsis* metaxin-like protein. Identical amino acids are 26% and similar amino acids are 42%. The identities are shown by the vertical red lines between the sequences, and the similarities by the red dots. *Arabidopsis* is included because it is the most widely studied model flowering plant. (B) Zebrafish metaxin 2 is aligned with the *Arabidopsis* metaxin-like protein. The identities are 21% and the similarities, 44%. In general, comparable percentages of identical amino acids are found in aligning vertebrate metaxin 1 and metaxin 2 proteins with a variety of plant metaxin-like proteins. (C) An invertebrate metaxin 1 sequence is aligned with a bacterial metaxin-like protein sequence. The invertebrate is *C. elegans*, an important model organism for molecular and developmental biology research. *Pseudomonas aeruginosa* is a common bacterium that is associated with hospital-acquired infections. The identities are 25% and the similarities, 43%, values that are close to the vertebrate metaxin 1 results in (A). (D) The alignment of *C. elegans* metaxin 2 with the *Pseudomonas* metaxin-like protein shows 23% identities and 43% similarities, values comparable to those in (B). The results with invertebrates and bacteria in (C) and (D) are typical of the results found for diverse invertebrate and bacterial species.

**Figure 3.**
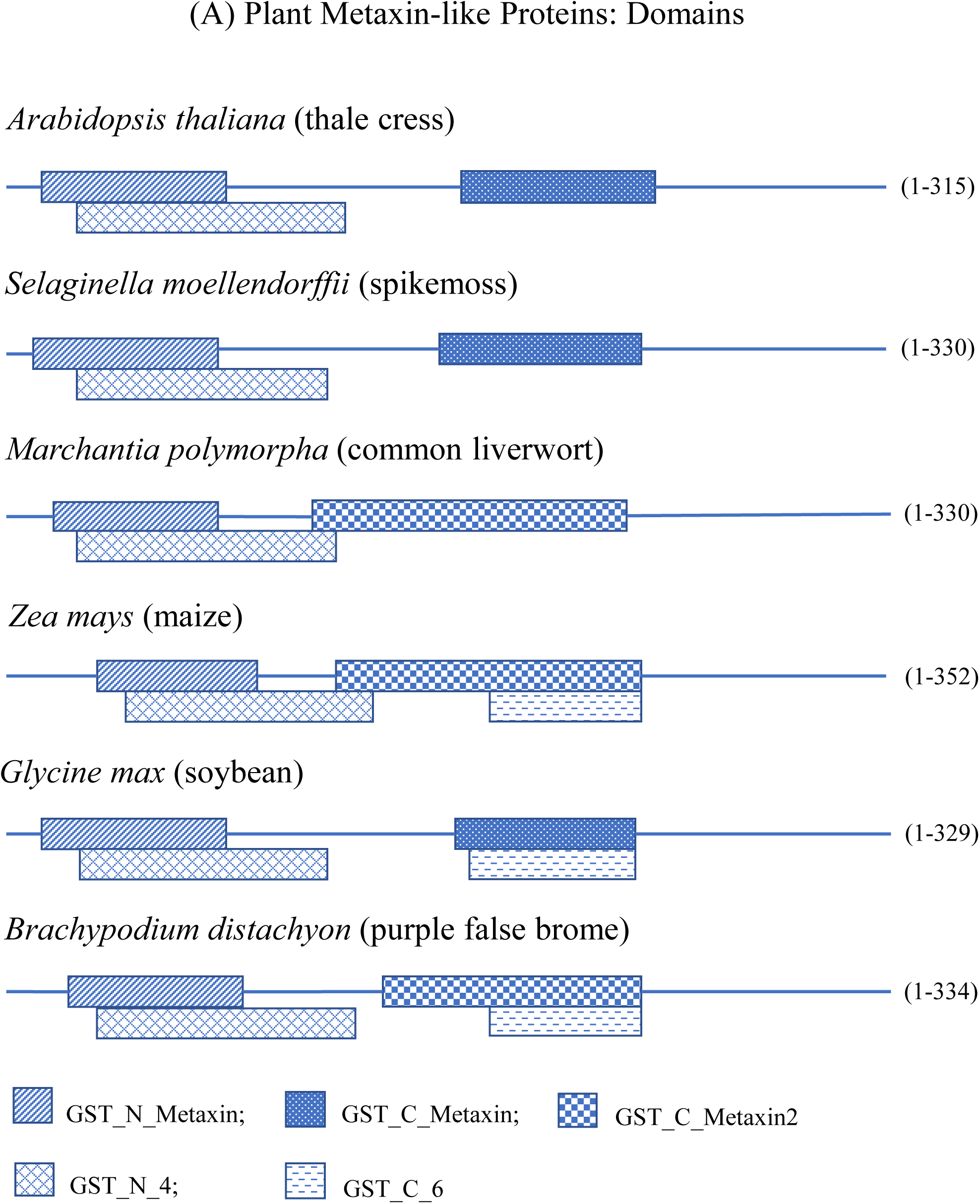
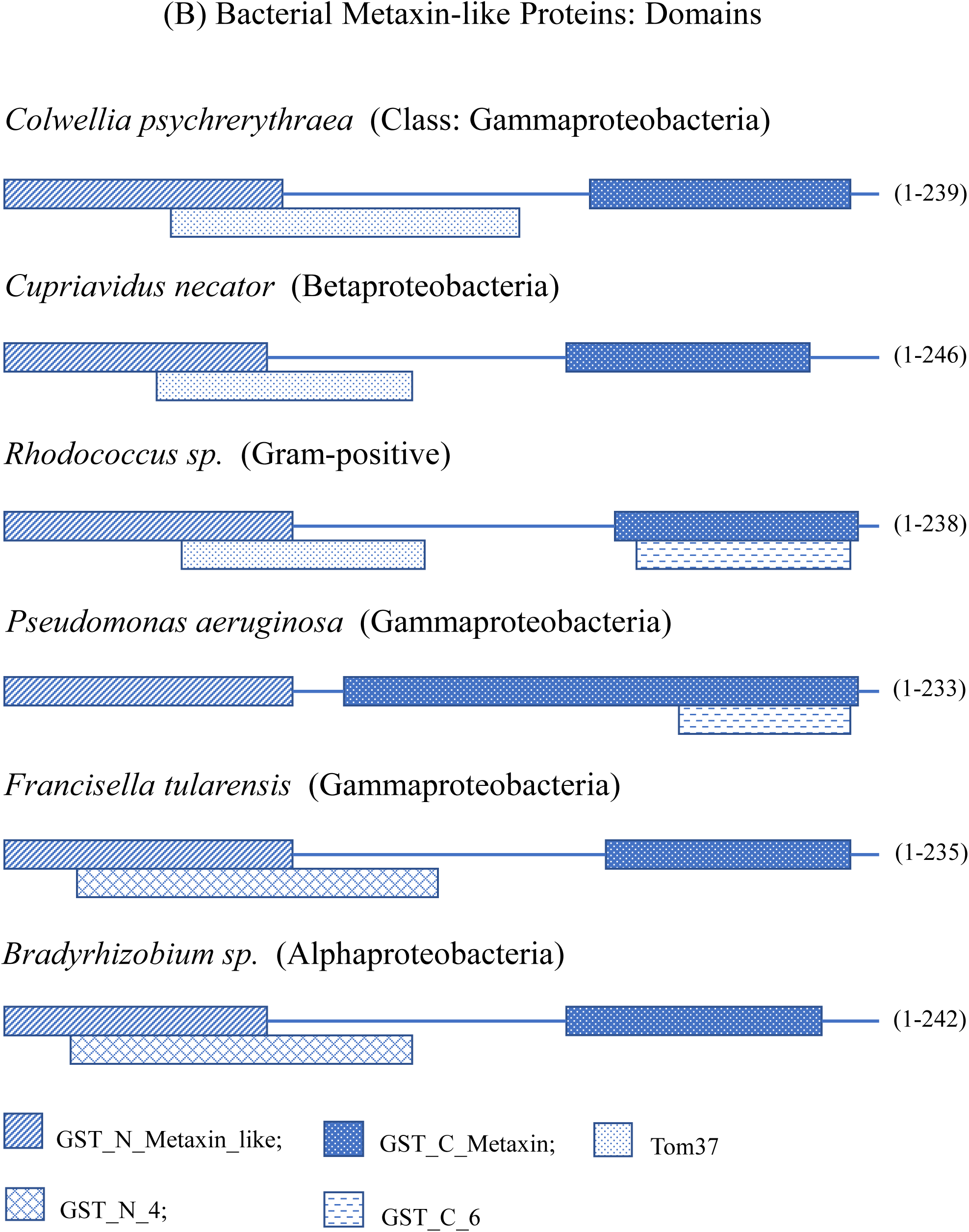
Domain structures of plant and bacterial metaxin-like proteins. (A) Conserved protein domains of representative plant metaxin-like proteins. The characteristic GST_N_Metaxin and GST_C_Metaxin domains are prominent domains in the figure. These domains are also a distinguishing feature of vertebrate and invertebrate metaxins. The figure shows the domains of eudicots, monocots, and non-angiosperms. The eudicots are *Arabidopsis* and the legume *Glycine max* (soybean). Maize or corn (*Zea mays*) and the experimental model grass, *Brachypodium distachyon*, are the monocots. Two plants are not angiosperms: *Selaginella moellendorffii* represents an ancient lineage of vascular plants, while the liverwort *Marchantia polymorpha* is a model non-vascular plant. In (B), the protein domain structures are shown for metaxin-like proteins from a variety of bacteria. As with the plant metaxin-like proteins, the bacterial proteins are characterized by the presence of GST_N_Metaxin and GST_C_Metaxin domains. Furthermore, the Tom37 domain, found for some of the bacterial proteins, is also a feature of the domain structures of vertebrate and invertebrate metaxins. The Gammaproteobacteria class, the most common taxonomic class of bacteria with metaxin-like proteins, is represented by *Colwellia, Pseudomonas*, and *Francisella. Bradyrhizobium* is an example of the Alphaproteobacteria, also a common bacterial class with metaxin-like proteins. *Cupriavidus* is in the Betaproteobacteria class. A bacterium that is not in the Proteobacteria phylum, *Rhodococcus* sp., is a Gram-positive bacterium of the phylum Actinobacteria.

**Figure 4.**
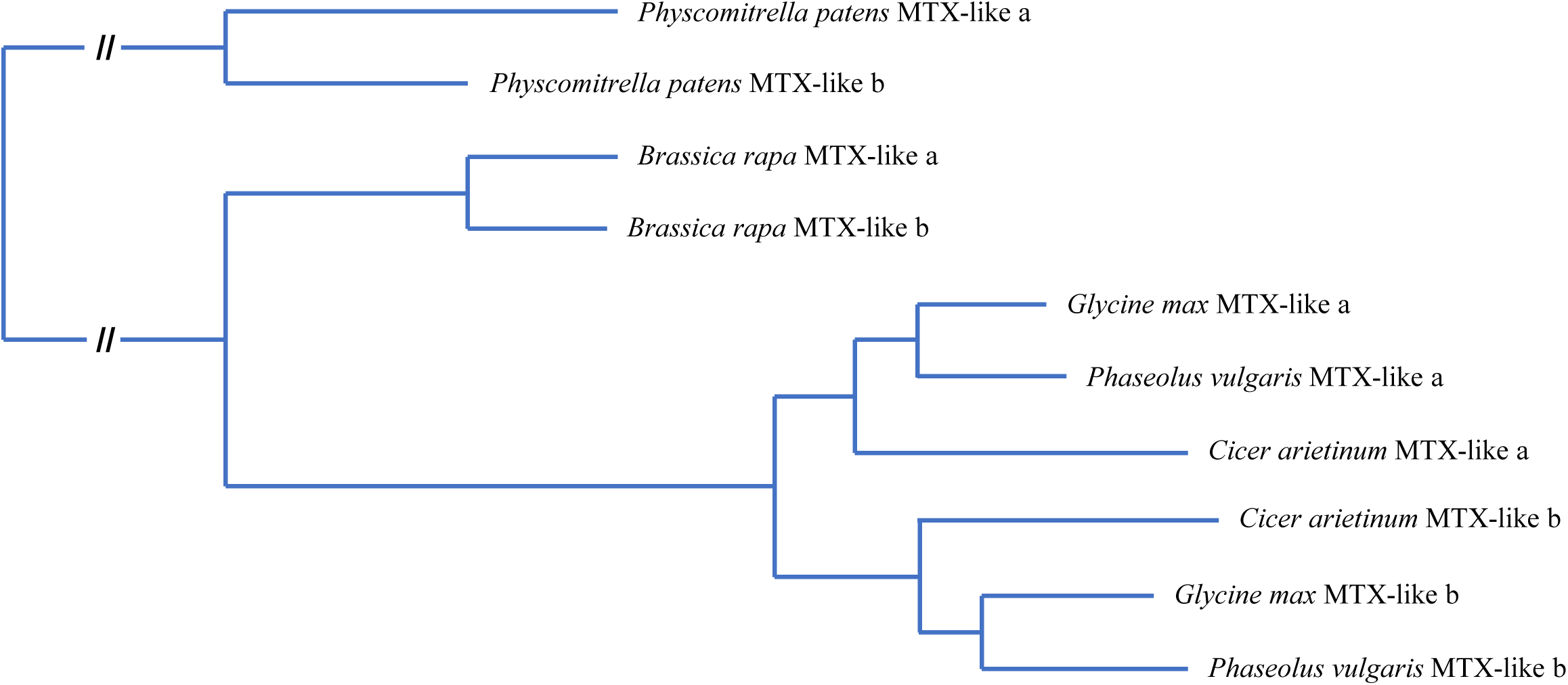
Detection of multiple metaxin-like proteins in plants. The figure shows a phylogenetic tree of representative metaxin-like proteins of plants that encode more than one protein. The two *Physcomitrella* metaxin-like a and b proteins group together, as do the two *Brassica* proteins. The *Glycine, Phaseolus*, and *Cicer* metaxin-like_a proteins form a group that is separate from the metaxin-like_b proteins. The two groups result from the greater homology of the metaxin-like proteins within a group, a or b, than between the a and b proteins of the same plant. The three plants in each group are in the same taxonomic family (Fabaceae), which is responsible for their high levels of amino acid sequence homology.

**Figure 5.**
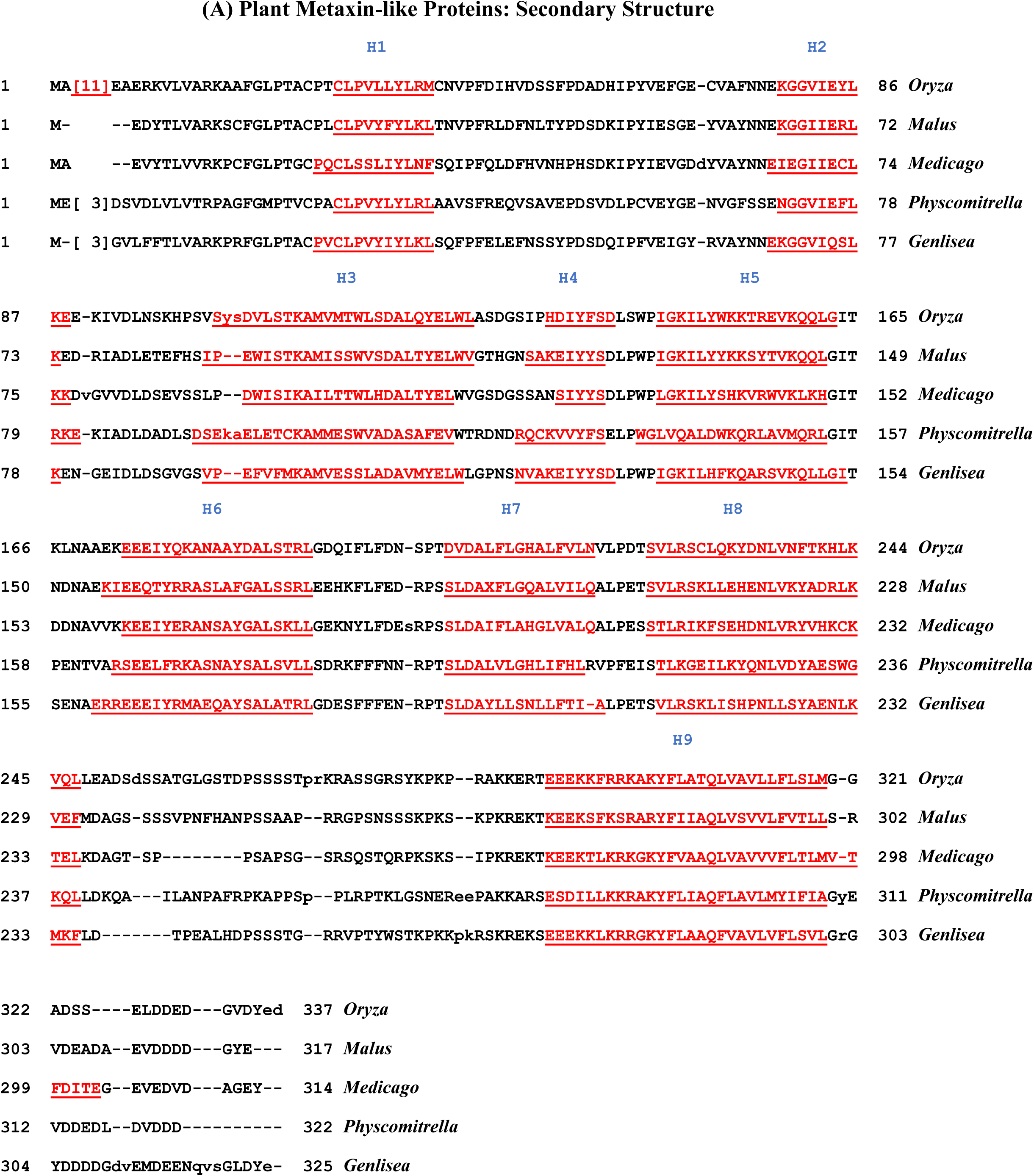
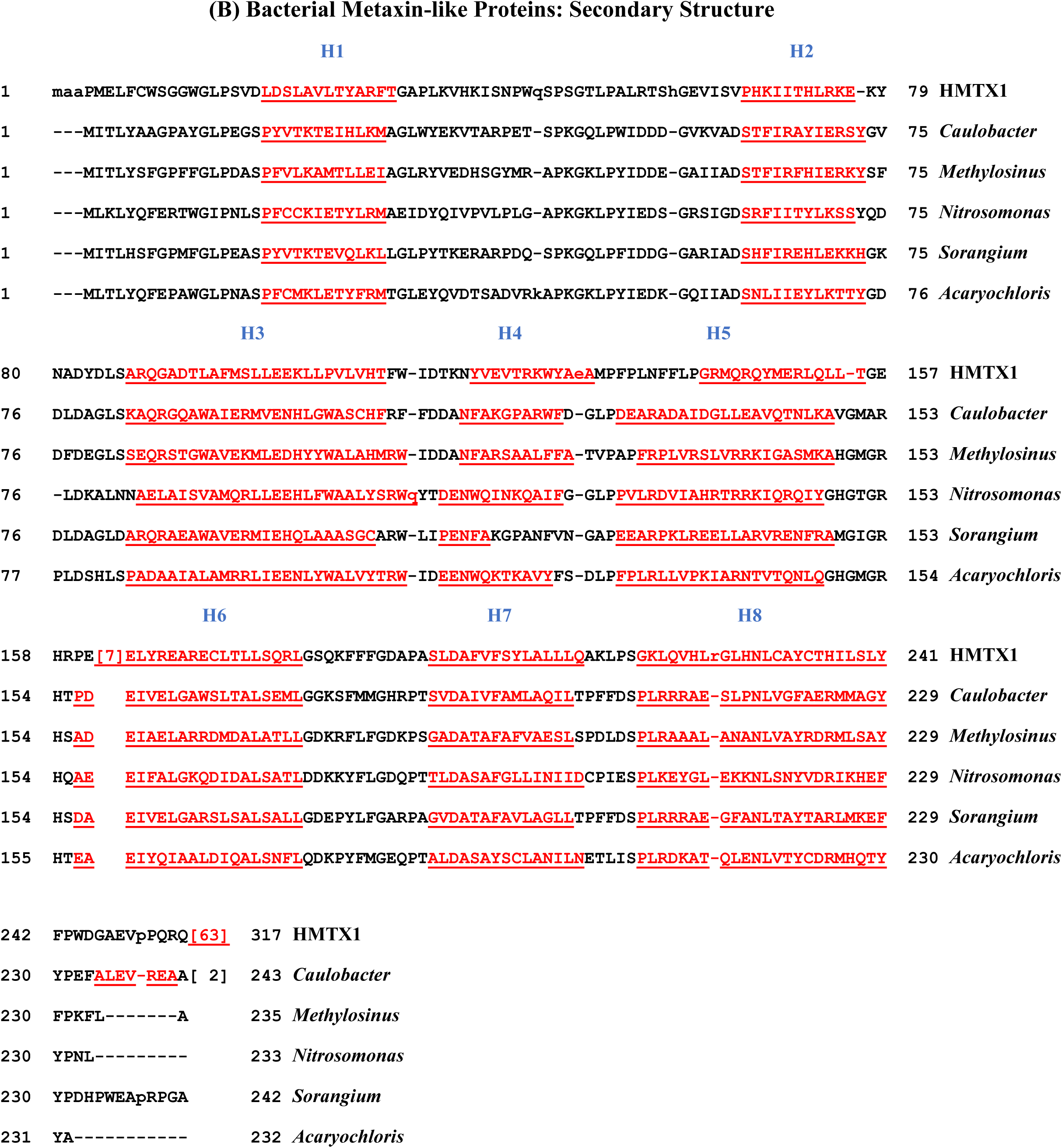
Multiple sequence alignments of alpha-helical segments of plant and bacterial metaxin-like proteins. In (A), the alpha-helical segments of plant metaxin-like proteins are underlined and shown in red. The conservation of helices, which are the predominant type of secondary structure, is a striking feature of these plant proteins. Beta-strand is largely absent. Both the location of the helices along the amino acid chains and the spacings between the segments are almost identical for each of the five plants included in the figure. The pattern of nine helices, H1 through H9, is the same as found for vertebrate metaxin 1 and 3 proteins. Vertebrate metaxin 2 lacks helix H9, but has an additional N-terminal helix. Three of the plants in the figure are model species: the legume *Medicago truncatula*, the moss *Physcomitrella patens*, and the carnivorous plant *Genlisea aurea*. The other two plants are important crop plants: Asian rice *Oryza sativa* and the apple *Malus domestica*. The arrangement of alpha-helical segments in bacterial metaxin-like proteins (Figure 5B) is generally similar to that of plants. A major difference, however, is the absence of helix H9 in the bacterial proteins. Vertebrate metaxin 2 proteins also lack H9. But, as discussed in section 3.3, bacterial metaxin-like proteins are about equally homologous to vertebrate metaxins 1, 2, and 3. Therefore, the bacterial proteins cannot be considered as prokaryotic metaxin 2 homologs, and the plant proteins as metaxin 1 or 3 homologs. The absence of helix 9 is reflected in the shorter lengths of the bacterial proteins at the C-terminal end. The shorter lengths can also be seen by comparing the protein domain structures of selected plant and bacterial metaxin-like proteins in Figure 3A and Figure 3B. In 3B, the GST_C_Metaxin domains of the bacterial proteins extend almost to the C-terminus of each.

## 3. RESULTS AND DISCUSSION

### 3.1. Existence of Metaxin-like Proteins in Plants and Bacteria

Among plants, metaxin-like proteins are found in a wide variety of angiosperms, both eudicots and monocots. The proteins, largely predicted from genomic sequences, were identified by their homology with vertebrate metaxins and the presence of characteristic GST_Metaxin protein domains. Metaxin-like proteins in eudicots are the most common. Examples of eudicots with metaxin-like proteins include the model flowering plant *Arabidopsis*, the potato, morning glory, and pumpkin. Among the monocots are wheat, the model grass *Brachypodium distachyon*, asparagus, and barley. Eudicots with metaxin-like proteins include both asterids and rosids. Examples of these asterids are *Genlisea aurea*, a carnivorous plant with the smallest known angiosperm genome, the tea plant, a wild tomato species, and a chili pepper. Rosids with metaxin-like proteins include the China rose, turnip, pink shepherd’s purse (a close relative of *Arabidopsis*), and the pomegranate. Other types of plants with metaxin-like proteins that are model plants in biology research are the moss *Physcomitrella patens* and the liverwort *Marchantia polymorpha*.

The bacterial species that have metaxin-like proteins are primarily in the Proteobacteria phylum. Gammaproteobacteria and Alphaproteobacteria are the most common taxonomic classes with these proteins. The Gammaproteobacteria in this study include species of *Pseudomonas, Francisella*, and *Colwellia*. For Gammaproteobacteria in general, *Legionella* species with metaxin-like proteins are particularly common. Among the Alphaproteobacteria in this study are *Caulobacter* and *Bradyrhizobium* species. Some Betaproteobacteria (example: *Cupriavidus*) and Deltaproteobacteria (*Sorangium*) also have metaxin-like proteins. In addition, other phyla have some representatives with these proteins: Cyanobacteria (example: *Acaryochloris*), Actinobacteria (*Rhodococcus*), and Nitrospirae (*Nitrospira*). Not all bacteria appear to have metaxin-like proteins. Searches of microbial protein databases have not revealed metaxin-like proteins for bacteria that include *Escherichia coli, Bacillus subtilis, Mycoplasma genitalium, Aliivibrio fischeri*, and *Synechocystis*.

### 3.2. Conservation of Plant and Bacterial Metaxin-like Proteins: Phylogenetic Analysis

As shown in Figure 1(A-1), phylogenetic analysis demonstrates that plant metaxin-like proteins, bacterial metaxin-like proteins, and vertebrate metaxins form discrete groups, but are related by evolution. The five selected examples of plants include *Arabidopsis thaliana* (a model flowering plant), *Populus trichocarpa* (the black cottonwood or poplar, a model tree species), *Glycine max* (soybean), *Medicago truncatula* (a model legume), and *Oryza sativa* (rice). Similar clustering, distinct from that of bacteria and vertebrates, was found for 39 other plants. Human, mouse, *Xenopus*, and zebrafish metaxin 1 and 3 proteins (MTX1 and MTX3) form the group of vertebrate metaxins in the figure. Vertebrate metaxin 2 proteins also form a cluster, not shown, that is separate but related to the other metaxins and metaxin-like proteins. The bacterial phyla in Figure 1(A-1) include Proteobacteria (*Pseudomonas, Parvibaculum, Colwellia*), Actinobacteria (*Rhodococcus*), Cyanobacteria (*Acaryochloris*), and Nitrospirae (*Nitrospira*). In Figure 1(A-2), phylogenetic analysis shows that plant metaxin-like proteins have evolutionary similarities that reflect similarities in the taxonomic classification of the plants. For these particular examples, *Arabidopsis* and *Tarenaya* differ at the level of taxonomic family, while the *Malus* and *Prunus* examples differ only at the level of genus. All four group together as eudicots. The *Phoenix* and *Elaeis* examples also differ only at the level of genus, and the two *Oryza* examples at the level of species. The latter four group together as monocots.

Figure 1B clearly reveals the difference between bacterial metaxin-like proteins and bacterial and human GSTs (glutathione S-transferase proteins). It shows that bacterial metaxin-like proteins are not bacterial GSTs. The group of bacterial metaxin-like proteins at the top of the figure is well separated from the bacterial and human GSTs at the bottom. In addition, the vertebrate metaxin 1 and metaxin 2 proteins of human, *Xenopus*, and zebrafish are seen to be more closely related to the bacterial metaxin-like proteins than to the GSTs. The bacterial GSTs in the figure are broadly representative of the GSTs found in bacteria. The variety of species with bacterial GSTs that are shown includes *E. coli, Salmonella, Yersinia, Burkholderia, Agrobacterium, Xanthomonas*, and *Actinobacillus*. The human GSTs also include a representative variety: GSTT1, GSTZ1, GSTM1, GSTP1, GSTA1, and GSTO1.

The observed grouping of the plant metaxin-like proteins in Figure 1A indicates that the genes encoding these proteins most likely arose from a common plant ancestor. Similarly, the separate grouping of the bacterial metaxin-like proteins in Figure 1(A-1) and Figure 1B suggests that their genes evolved from the same ancestral bacterial gene. Further, the phylogenetic analysis reveals that the plant, bacterial, and vertebrate proteins are all related by evolution. It therefore appears that the genes for the metaxin-like proteins and vertebrate metaxins originated from the same ancestral gene. Detailed analysis shows that the plant and bacterial metaxin-like proteins are about equally homologous to vertebrate metaxins 1, 2, and 3. Which vertebrate metaxin gene is most directly related to the common ancestral gene will require further investigation.

### 3.3. Amino Acid Sequence Alignments: Homology of the Metaxin-like Proteins

Plant metaxin-like proteins show a similar degree of homology to the three vertebrate metaxins: 1, 2, and 3. For example, the *Arabidopsis thaliana* metaxin-like protein (315 amino acids) is 26% identical to human metaxin 1 (317 aa), 22% identical to human metaxin 2 (263 aa), and 28% identical to human metaxin 3 (326 aa). For the *Arabidopsis* metaxin-like protein and zebrafish (*Danio rerio*) metaxins 1a and 2, the numbers are 26% and 21% (Figure 2A,B), and for zebrafish metaxin 3, 31%. Zebrafish metaxin 1a is one of two zebrafish metaxin 1 proteins, 1a and 1b. Comparable results are found for a variety of plant metaxin-like proteins, demonstrating that these proteins are about equally homologous to all three vertebrate metaxins. There is no significant homology, however, between the plant metaxin-like proteins and human GSTs. Similar percentages of identities are found comparing plant metaxin-like proteins with bacterial metaxin-like proteins. As examples, the metaxin-like proteins of the bacteria *Rhodococcus, Parvibaculum, Nitrospira*, and *Colwellia* have an average of 24% identities relative to the *Arabidopsis* metaxin-like protein. Higher percentages, around 50%, are found in comparing one plant metaxin-like protein with another. *Arabidopsis* and *Glycine max* (soybean) have 51% identities, *Hordeum vulgare* (barley) and *Medicago truncatula* (a model legume) have 47%, *Oryza sativa* (rice) and *Populus trichocarpa* (poplar) have 52%, while *Selaginella moellendorffii* (a model organism of evolutionary biology) and *Sorghum bicolor* (sorghum) have 39%.

Bacterial metaxin-like proteins, like the plant proteins, display similar degrees of homology to vertebrate metaxins 1, 2, and 3. An example is the *Pseudomonas aeruginosa* metaxin-like protein (233 amino acids), which has 28%, 23%, and 22% identical amino acids compared to human metaxins 1, 2, and 3, respectively. When aligned with the *Pseudomonas* metaxin-like protein, metaxins 1, 2, and 3 of another vertebrate, *Xenopus laevis*, show 22%, 21%, and 24% identities, respectively. In Figure 2C,D, metaxins 1 and 2 of the model invertebrate *C. elegans* are aligned with the *Pseudomonas* metaxin-like protein: 25% identities (metaxin 1) and 23% identities (metaxin 2) are found. As mentioned above, aligning various bacterial metaxin-like proteins with plant metaxin-like proteins shows similar degrees of homology. To give an additional example, the metaxin-like protein of the bacterium *Francisella tularensis* aligned with the metaxin-like proteins of the plants *Phaseolus vulgaris* (bean), *Panicum virgatum* (switchgrass), *Vitis vinifera* (grape vine), *Sorghum bicolor* (sorghum), and *Setaria italica* (millet) reveals amino acid identities of 26%, 24%, 26%, 24%, and 26%, respectively. Comparing one bacterial metaxin-like protein with another shows about 38% amino acid identities. As examples, *Aquicella lusitana* and *Thiotrichaceae* sp. have 39% identical amino acids, *Oceanospirillum* sp. and *Solimonas fluminis* have 37%, *Cycloclasticus* sp. and *Zooshikella ganghwensis* have 38%, and *Alphaproteobacteria* sp. and *Sandaracinus amylolyticus* have 39%.

Table 1 includes fundamental properties of the predicted amino acid sequences of selected plant and bacterial metaxin-like proteins. The numbers of amino acids and molecular weights of the plant metaxin-like proteins are similar to those of vertebrate metaxins 1 and 3, in particular human metaxin 1 (317 amino acids) and metaxin 3 (326 aa). The bacterial metaxin-like proteins have shorter protein chains, closer to that of human metaxin 2 (263 aa). The difference is due to the extended C-terminal sequences of the plant metaxin-like proteins. This can be seen in comparing Figure 3A with 3B. The figure, showing the protein domains of the metaxin-like proteins, is discussed in more detail in Section 3.4. The extended C-terminal regions of the plant metaxin-like proteins lack any additional protein domains. But the C-terminal regions do contain an extra alpha-helical segment not found with the bacterial proteins and designated as helix H9 in Figure 5A.

**Table 1.**
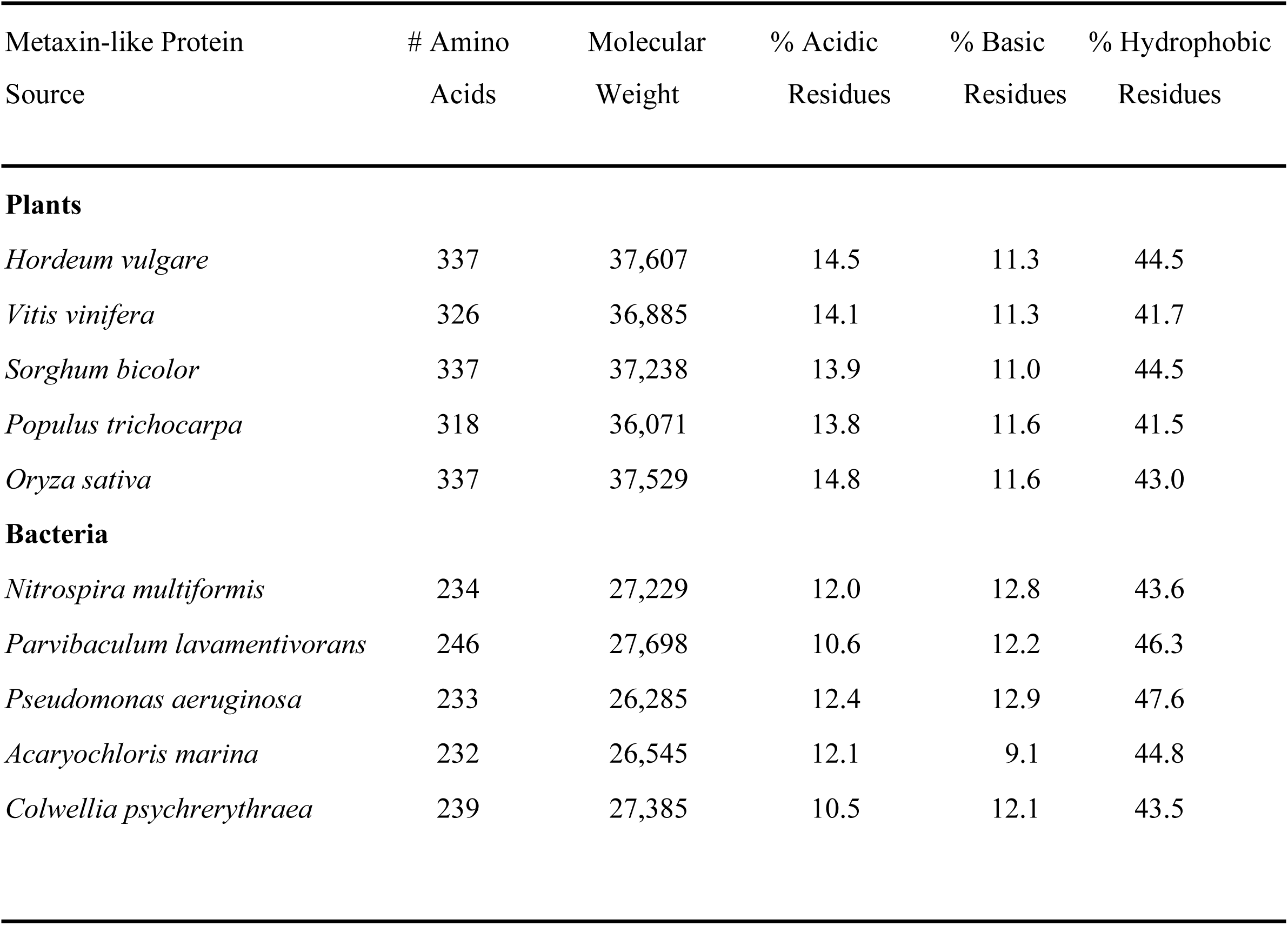
Amino Acid Analysis of Plant and Bacterial Metaxin-like Proteins.

### 3.4. Protein Domains of Plant and Bacterial Metaxin-like Proteins

Metaxin proteins are characterized by the presence of GST_N_Metaxin and GST_C_Metaxin conserved domains. This is true for vertebrate metaxins 1, 2, and 3, invertebrate metaxins 1 and 2, and, as Figure 3 demonstrates, the plant and bacterial metaxin-like proteins. In Figure 3A, both model plants (*Arabidopsis, Selaginella, Marchantia, Brachypodium*) and economically important crop plants (maize or corn, soybean) are shown. Besides the presence of GST_Metaxin domains, the lengths of the protein chains are comparable to the lengths that characterize vertebrate metaxins 1 and 3, such as human metaxins 1 and 3 with 317 and 326 amino acids, respectively. Three of the plant proteins in Figure 3 possess GST_C_Metaxin2 domains also found with metaxin 2 proteins. However, as discussed in section 3.3, both the plant and bacterial metaxin-like proteins are about equally homologous to vertebrate metaxins 1, 2, and 3. The plant metaxin-like proteins cannot therefore be subdivided into metaxin 1-like and metaxin 2-like proteins.

As with the plant metaxin-like proteins, the bacterial proteins are also characterized by GST_N_Metaxin and GST_C_Metaxin domains (Figure 3B). The presence of a Tom37 domain in some bacterial proteins (*Colwellia, Cupriavidus, Rhodococcus* in 3B) is also a feature of vertebrate and invertebrate metaxins. The number of amino acids in the bacterial metaxin-like proteins is less than in the plant metaxin-like proteins. The examples in Figure 3B show that these bacterial proteins have a similar length to human metaxin 2 and other vertebrate metaxin 2 proteins. As mentioned in section 3.3 in connection with Table 1, the difference with the plant metaxin-like proteins is largely due to the shorter C-terminal end of the bacterial proteins. This results in the GST_C_Metaxin domain being close to the C-terminus.

In addition to metaxin-like proteins with GST_Metaxin domains, bacteria also contain GST-like proteins with different GST domains. And individual bacteria can contain a variety of GST-like proteins. For example, database searches reveal that *Pseudomonas aeruginosa* (strain PA01) contains GST-like proteins with GST_N_Beta and GST_C_Beta domains, GST_N_Zeta and GST_C_Zeta domains, GST_N_2 and GST_C_1 domains, and GST_N_Ure2p_like and GST_C_Ure2p_like domains. Additional proteins of the GST type also exist in various bacteria. *Legionella drancourtii*, for example, has only one GST-like protein, with GST_N_SspA and GST_C_SspA domains.

### 3.5. Presence of Multiple Metaxin-like Proteins in Plants

Multiple metaxin-like proteins were found for about a quarter of the different plant species studied. The phylogenetic relationships of five examples are shown in Figure 4. The examples include *Physcomitrella patens* (a model moss), *Brassica rapa* (turnip, bok choy, field mustard), *Glycine max* (soybean), *Phaseolus vulgaris* (common bean), and *Cicer arietinum* (chickpea). Two metaxin-like proteins, a and b, were detected in each case, except for *Glycine max*. The two proteins of these examples show a high level of homology. In particular, the two *Physcomitrella* metaxin-like proteins have 72% amino acid identities, the two *Brassica* proteins 86%, the *Phaseolus* proteins 69%, and the *Cicer* proteins 63%. For *Glycine max*, four metaxin-like proteins were detected, a, b, c, d, all on different chromosomes (chromosomes 9, 13, 15, and 17, respectively). The figure shows two of these, a and b. The metaxin-like_a protein (329 amino acids) shares 71% amino acid sequence identities with metaxin-like_b (320 aa), 91% with metaxin-like_c (328 aa), and 70% with metaxin-like_d (321 aa). But these are not homologs of metaxins 1, 2, and 3 found in vertebrates. The percentages of amino acid identities for *Glycine max* metaxin-like_a aligned with human metaxins 1, 2, and 3 are 27%, 24%, and 20%, respectively, while the comparable percentages for *Glycine max* metaxin-like_b are 21%, 24%, and 23%. Similar percentages are found for c and d. Therefore, the soybean metaxins are about equally homologous to all three human metaxins. The same is true for the plants with two metaxin-like proteins. As an example, the *Physcomitrella* metaxin-like_a protein has 24%, 26%, and 23% amino acid identities when aligned with human metaxins 1, 2, and 3, respectively. For metaxin-like_b, the percentages are 23%, 28%, and 24%.

Figure 4 indicates that the metaxin-like_a and metaxin-like_b proteins of *Glycine, Phaseolus*, and *Cicer* form separate, distinct a and b groups. This is due to the three metaxin-like proteins in each of the two groups, a and b, being from plants of the same taxonomic family (Fabaceae). The result is a higher percentage of amino acid identities within a group than for the a and b proteins of the same plant. For example, the *Glycine max* a and b proteins align with 71% identities, but Glycine *max*_a has 87% identities aligned with *Phaseolus vulgaris*_a, and 75% with *Cicer arietinum*_a.

### 3.6. Neighboring Genes of Metaxin-like Protein Genes

The genes adjacent to the plant metaxin-like protein genes show only a weak degree of conservation in comparing one plant with another, with more differences than similarities. For example, comparing the genomic regions next to the *Glycine max* metaxin-like_a gene and the *Zea mays* metaxin-like gene shows just 1 out of 21 *Zea mays* genes to be identical to the *Glycine max* genes. Comparing the *Glycine max* metaxin-like_a genomic region with that of the model plant *Arabidopsis* shows 0 out of 19 *Arabidopsis* genes to be identical to the *Glycine max* genes. Although only weak conservation is the general rule, a higher degree of conservation of neighboring genes may be found for some plants. As examples, four genes that are adjacent to the *Glycine max* metaxin-like_a gene are also neighboring genes of the metaxin-like proteins of *Cicer arietinum*_a, *Citrus sinensis* (sweet orange), and *Phoenix dactylifera* (date palm). These neighboring genes are in the order: **metaxin-like protein**/pumilio homolog/U3 small nucleolar RNA-associated protein/villin-4-like/putative glucose-6-phosphate 1-epimerase.

Comparison of the genes adjacent to the genes of selected bacterial metaxin-like proteins shows that the genomic context is not conserved. The selected bacteria included *Pseudomonas, Colwellia, Acaryochloris, Nitrosomonas*, and *Bradyrhizobium*. For example, identified genes that are next to the gene for the *Pseudomonas* metaxin-like protein are in the order: carboxylesterase/amino acid binding protein/phosphatidylcholine synthase/**metaxin-like protein**/GIY-YIG nuclease superfamily/transferase. But in *Colwellia*, the genes are: inosine 5’monophosphate dehydrogenase/GMP synthase/L-PSP family endoribonuclease/**metaxin-like protein**/RecQ domain-containing protein/hemerythrin family protein/membrane protein.

### 3.7. Alpha-helical Secondary Structures of Plant and Bacterial Metaxin-like Proteins

As demonstrated by Figure 5, the secondary structures of the metaxin-like proteins are dominated by a highly conserved pattern of alpha-helices. The five representative plant metaxin-like proteins in Figure 5A show the same pattern of nine helical segments, H1 to H9. Furthermore, the spacings between pairs of helices are almost identical for each of the five examples. The pattern of nine alpha-helices is also found for vertebrate metaxins 1 and 3. The plants included in the figure are *Oryza sativa* (rice), *Malus domestica* (apple), *Medicago truncatula* (a model legume), *Physcomitrella patens* (a model moss), and *Genlisea aurea* (a carnivorous plant with an extremely small genome).

For bacterial metaxin-like proteins (Figure 5B), a highly conserved pattern of alpha-helices is also found. The five bacterial protein sequences in the figure have the same eight helices, H1 to H8, with similar spacings between pairs of helices for the five examples. These helices are identical to helices H1 to H8 of human metaxin 1 (HMTX1), also included in the figure. Human metaxin 1 has, in addition, a ninth helix, H9, that is C-terminal and in the sequence denoted by [63] at the C-terminus. Other vertebrate metaxin 1 proteins are similar. Moreover, the bacterial metaxin-like proteins don’t have the extra N-terminal helix found for vertebrate metaxin 2, including human metaxin 2. They therefore lack both the extra C-terminal helix of vertebrate metaxin 1 (helix H9) and the extra N-terminal helix of vertebrate metaxin 2. In general, though, the conservation of alpha-helices is a prominent feature of the secondary structures of metaxins and metaxin-like proteins of many different organisms: plants, bacteria, vertebrates, invertebrates. The bacteria in Figure 5B are *Caulobacter* sp., *Methylosinus trichosporium, Nitrosomonas* sp., *Sorangium cellulosum*, and *Acaryochloris marina*.

A transmembrane alpha-helical segment near the C-terminus is a feature of the metaxin 1 proteins of vertebrates and invertebrates. In this study, plant metaxin-like proteins were also found to have C-terminal transmembrane helices. For example, the *Arabidopsis thaliana* metaxin-like protein is predicted to have a transmembrane helix between amino acids 284 and 303 of the 315 residue protein. *Zea mays* (maize or corn) has a helix between amino acids 258 and 277 of the 292 residue protein. The transmembrane helices of these examples overlap with the C-terminal 2/3 of helix H9 in Figure 5A. Transmembrane helices appear to be a general feature of plant metaxin-like proteins and were found for all of the proteins investigated. However, bacterial metaxin-like proteins were not found to have transmembrane helices. This seems related to the shorter lengths of the bacterial metaxin-like proteins as seen in Figure 3B and the absence of helix H9.

## REFERENCES

Adolph, K.W. (2004). The zebrafish metaxin 3 gene (*mtx3*): cDNA and protein structure, and comparison to zebrafish metaxins 1 and 2. Gene 330, 67–73.

Adolph, K.W. (2005). Characterization of the cDNA and amino acid sequences of *Xenopus* metaxin 3, and relationship to *Xenopus* metaxins 1 and 2. DNA Sequence 16, 252–259.

Adolph, K.W. (2019). Metaxin 3 is a highly conserved vertebrate protein homologous to mitochondrial import proteins and GSTs. bioRxiv doi: 10.1101/813451. (http://biorxiv.org/cgi/content/short/813451v1)

Adolph, K.W. (2020). Invertebrate metaxins 1 and 2: widely distributed proteins homologous to vertebrate metaxins implicated in protein import into mitochondria. bioRxiv doi: 10.1101/2020.01.06.895979. (https://www.biorxiv.org/content/10.1101/2020.01.06.895979v1)

Adolph, K.W., Long, G.L., Winfield, S., Ginns, E.I., and Bornstein, P. (1995). Structure and organization of the human thrombospondin 3 gene (*THBS3*). Genomics 27, 329–336.

Altschul, S.F., Madden, T.L., Schaeffer, A.A., Zhang, J., Zhang, Z., Miller, W., and Lipman, D.J. (1997). Gapped BLAST and PSI-BLAST: a new generation of protein database search programs. Nucleic Acids Res. 25, 3389–3402.

The *Arabidopsis* Genome Initiative (2000). Analysis of the genome sequence of the flowering plant *Arabidopsis thaliana*. Nature 408, 796–815.

Armstrong, L.C., Komiya, T., Bergman, B.E., Mihara, K., and Bornstein, P. (1997). Metaxin is a component of a preprotein import complex in the outer membrane of the mammalian mitochondrion. J. Biol. Chem. 272, 6510–6518.

Armstrong, L.C., Saenz, A.J., and Bornstein, P. (1999). Metaxin 1 interacts with metaxin 2, a novel related protein associated with the mammalian mitochondrial outer membrane. J. Cell. Biochem. 74, 11–22.

The French-Italian Public Consortium for Grapevine Genome Characterization (2007). The grapevine genome sequence suggests ancestral hexaploidization in major angiosperm phyla. Nature 449, 463–467.

Jones, D.T. (1999). Protein secondary structure prediction based on position-specific scoring matrices. J. Mol. Biol. 292, 195–202.

Krogh, A., Larsson, B., von Heijne, G., and Sonnhammer, E.L.L. (2001). Predicting transmembrane protein topology with a hidden Markov model: application to complete genomes. J. Mol. Biol. 305, 567–580.

Larsson, P., Oyston, P., Chain, P. et al. (2005). The complete genome sequence of *Francisella tularensis*, the causative agent of tularemia. Nat. Genet. 37, 153–159.

Long, G.L., Winfield, S., Adolph, K.W., Ginns, E.I., and Bornstein, P. (1996). Structure and organization of the human metaxin gene (*MTX*) and pseudogene. Genomics 33, 177–184.

Lucker, S., Wagner, M., Maixner, F. et al. (2010). A *Nitrospira* metagenome illuminates the physiology and evolution of globally important nitrite-oxidizing bacteria. Proc. Natl. Acad. Sci. U.S.A. 107, 13479–13484.

Madeira, F., Park, Y.M., Lee, J., Buso, N., Gur, T., Madhusoodanan, N., Basutkar, P., Tivey, A.R.N., Potter, S.C., Finn, R.D., and Lopez, R. (2019). The EMBL-EBI search and sequence analysis tools APIs in 2019. Nucleic Acids Res. 47, W636–W641.

Marchler-Bauer, A., Bo, Y., Han, L., He, J., Lanczcki, C.J., Lu, S., Chitsaz, F., Derbyshire, M.K., Geer, R.C., Gonzales, N.R., Gwadz, M., Hurwitz, D.I., Lu, F., Marchler, G.H., Song, J.S., Thanki, N., Wang, Z., Yamashita, R.A., Zhang, D., Zheng, C., Geer, L.Y., and Bryant, S.H. (2017). CDD/SPARCLE: functional classification of proteins via subfamily domain architectures. Nucleic Acids Res. 45, D200–D203.

Methe, B.A., Nelson, K.E., Deming, J.W. et al. (2005). The psychrophilic lifestyle as revealed by the genome sequence of *Colwellia psychrerythraea* 34H through genomic and proteomic analyses. Proc. Natl. Acad. Sci. U.S.A. 102, 10913–10918.

Needleman, S.B. and Wunsch, C.D. (1970). A general method applicable to the search for similarities in the amino acid sequences of two proteins. J. Mol. Biol. 48, 443–453.

Neupert, W. (2015). A perspective on transport of proteins into mitochondria: a myriad of open questions. J. Mol. Biol. 427, 1135–1158.

Ouyang, S., Zhu, W., Hamilton, J. et al. (2007). The TIGR Rice Genome Annotation Resource: improvements and new features. Nucleic Acids Res. 35, D883–D887.

Papadopoulos, J.S. and Agarwala, R. (2007). COBALT: constraint-based alignment tool for multiple protein sequences. Bioinformatics 23, 1073–1079.

Pfanner, N., Warscheid, B., and Wiedemann, N. (2019). Mitochondrial proteins: from biogenesis to functional networks. Nat. Rev. Mol. Cell Biol. 20, 267–284.

Rensing, S.A, Lang, D., Zimmer, A.D. et al. (2008). The *Physcomitrella* genome reveals evolutionary insights into the conquest of land by plants. Science 319, 64–69.

Stover, C.K., Pham, X.Q., Erwin, A.L. et al. (2000). Complete genome sequence of *Pseudomonas aeruginosa* PA01, an opportunistic pathogen. Nature 406, 959–964.

Swingley, W.D., Chen, M., Cheung, P.C. et al. (2008). Niche adaptation and genome expansion in the chlorophyll d-producing cyanobacterium *Acaryochloris marina*. Proc. Natl. Acad. Sci. U.S.A. 105, 2005–2010.

Tuskan, G.A., Difazio, S., Jansson, S. et al. (2006). The genome of black cottonwood, *Populus trichocarpa* (Torr. & Gray). Science 313, 1596–1604.

Wiedemann, N. and Pfanner, N. (2017). Mitochondrial machineries for protein import and assembly. Annu. Rev. Biochem. 86, 685–714.

